# Genetic or toxicant induced disruption of vesicular monoamine storage and global metabolic profiling in *Caenorhabditis elegans*

**DOI:** 10.1101/2020.10.02.324095

**Authors:** Joshua M. Bradner, Vrinda Kalia, Fion K. Lau, Monica Sharma, Meghan L. Bucher, Michelle Johnson, Merry Chen, Douglas I. Walker, Dean P. Jones, Gary W. Miller

## Abstract

The proper storage and release of monoamines contributes to a wide range of neuronal activity. Here, we examine the effects of altered vesicular monoamine transport in the nematode *C. elegans*. The gene *cat-1* is responsible for the encoding of the vesicular monoamine transporter (VMAT) in *C. elegans* and is analogous to the mammalian vesicular monoamine transporter 2 (VMAT2). Our laboratory has previously shown that reduced VMAT2 activity confers vulnerability on catecholamine neurons in mice. The purpose of this paper was to determine whether this function is conserved and to determine the impact of reduced VMAT activity in *C. elegans*. Here we show that deletion of *cat-1*/VMAT increases sensitivity to the neurotoxicant 1-methyl-4-phenylpyridinium (MPP^+^) as measured by enhanced degeneration of dopamine neurons. Reduced *cat-1*/VMAT also induces changes in dopamine-mediated behaviors. High-resolution mass spectrometry-based metabolomics in the whole organism reveals changes in amino acid metabolism, including tyrosine metabolism in the *cat-1*/VMAT mutants. Treatment with MPP^+^ disrupted tryptophan metabolism. Both conditions altered glycerophospholipid metabolism, suggesting a convergent pathway of neuronal dysfunction. Our results demonstrate the evolutionarily conserved nature of monoamine function in *C. elegans* and further suggest that HRMS-based metabolomics can be used in this model to study environmental and genetic contributors to complex human disease

## Introduction

The synaptic vesicle plays a significant role in the protection of neurons from toxic insult. The vesicular amine transporters (VATs) are members of the toxin extruding antiporter (TEXAN) gene family of proton antiporters, closely related to proteins in prokaryotic organisms that work to exclude antibiotics from the cytoplasm (Parsons 2000; Schuldiner et al. 1995). In eukaryotes, members of this family reside on the inner vesicular membrane and are instrumental in packaging neurotransmitters into synaptic vesicles, ensuring the transmission of the action potential. Additionally, these transporters have cytoprotective effects, keeping potentially toxic by-products of neurotransmitters out of the cytoplasm (vesicular monoamine transporter 2, VMAT2) and ensuring the replenishment of amine stores of acetylcholine in the synaptic cleft (vesicular acetylcholine transporter, VAChT). The breakdown in this relationship can have potentially deadly results. The adequate release of acetylcholine into the synapse is dependent on the vesicular acetylcholine transporter (VAChT). Impairment of acetylcholine release into the synaptic cleft, either through direct inhibition of VAChT (using vesamicol) or indirectly (using botulinum toxin) can lead to respiratory failure and death (Burton et al. 1994; Dressler et al. 2005; Prior et al. 1992). For other members of the TEXAN family, the breakdown of antiporter activity in synaptic vesicles appears to result in direct toxicity to the cell from oxidized amines, like oxidized metabolites of dopamine. (Spina and Cohen 1989).

When its storage is dysregulated, dopamine can accumulate in the cytosol where it is vulnerable to oxidation and enzymatic catabolism. These processes generate reactive metabolites, such as the dopamine quinone, as well as oxidative stress inducing reactive oxygen species (Stokes et al. 1999) and catechol aldehydes (Goldstein et al. 2013; Vermeer et al. 2012). Through the utility of mouse, *in-vitro* vesicular uptake, and cell culture models, we have established considerable evidence that VMAT2 confers resistance to toxic insult in dopaminergic neurons as a graded response in vesicular function (Caudle et al. 2007; Fumagalli et al. 1999; Lohr et al. 2014; Lohr et al. 2016; Lohr et al. 2015). We have proposed this increase in VMAT2 leads to a proportional increase in the storage capacity of monoaminergic synaptic vesicles (Lohr et al. 2014). This in turn provides increased protection from both 1-methyl-4-phenyl-1,2,3,6-tetrahydropyridine (MPTP) and N-methylamphetamine (METH) toxicity (Lohr et al. 2014; Lohr et al. 2016; Lohr et al. 2015). Others have demonstrated similar results including VMAT2-mediated protection against L-DOPA (the precursor to dopamine) and Parkinson’s disease associated toxicants (Lawal et al. 2010; Mosharov et al. 2009; Munoz et al. 2012). The hypothesis that increased vesicular sequestration capacity is able to protect against endogenous and exogenous toxicants has been supported in human studies that show a reduction in Parkinson’s disease risk associated with gain of function haplotypes in the promoter region of VMAT2 (Brighina et al. 2013; Glatt et al. 2006).

Recent findings highlight VMAT2 as a vulnerable target to environmental toxicants (Caudle 2015; Caudle et al. 2012). Animal studies suggest that a range of chemical exposures including pesticides, fungicides, polychlorinated biphenyls (PCBs), polybrominated diphenyl ethers (PBDE’s), and perfluorinated compounds reduce the expression and function of VMAT2 and lead to symptoms closely resembling Parkinson’s Disease (PD) (Bradner et al. 2013; Caudle et al. 2005; Choi et al. 2015; Enayah et al. 2018; Inamdar et al. 2013; Miller et al. 1999; Patel et al. 2016; Pham-Lake et al. 2017; Richardson et al. 2006; Richardson et al. 2008; Richardson and Miller 2004; Schuh et al. 2009; Wilson et al. 2014; Xiong et al. 2016). Continued research into the effects of these chemicals on the function of amine transporters should improve our understanding of how they function to disrupt the vesicular integrity of the synaptic vesicle.

Despite the notable benefits of mouse and *in vitro* cell culture models, changes in behavior and alterations in whole organism metabolic function are difficult and costly to perform in mammalian models. As such, our lab has begun using *C. elegans* as a method to measure neurotoxic outcomes as mediated by a protein 49% identical to VMAT2, a monoamine transporter encoded by the gene *cat-1* (Duerr et al. 1999). In this manuscript we evaluate the effects of the *cat-1* (*ok411*) mutant allele on *cat-1*/VMAT expression, dopamine neuron integrity, and monoamine-dependent behaviors. We also determined the effects of reduced *cat-1*/VMAT on vulnerability to the classical Parkinson’s disease-associated toxicant 1-methyl-4-phenylpyridinium (MPP^+^). Finally, we use high-resolution mass spectrometry-based metabolomics to determine the impact of cat-1/VMAT reduction or MPP^+^ treatment on metabolism, measured using the whole organism.

## Materials and Methods

### Reagents

Except where noted, all reagents were purchased from Sigma Aldrich (St. Louis, MO).

### Phylogenetic relationships of vesicular amine transporters

Sequences of vesicular amine transporters (supplemental data) from various species were subjected to cladistic organization via bioinformatics online tools locate at www.phylogeny.fr (Dereeper et al. 2008).

### Strains and Culture Conditions

Long established standard methods of culture including the use of normal growth media (NGM) plates, culture temperatures of 20° C and the OP50 *E. coli* strain as a food source were followed as described (Brenner 1974). Deviations in this practice are mentioned in conjunction with the relevant assay. *C. elegans* strains were provided by the CGC, which is funded by NIH Office of Research Infrastructure Programs (P40 OD010440). These include the wild type N2 Bristol strain, *dat-1* (*ok157*) mutant strain, RB681 [*cat-1*(*ok411*)] mutant strain, the fluorescent transgenic strain BZ555 (*pDAT*::GEP), and a cross between strain RB681 and BZ555 [(*pDAT*::GFP; *cat-1*(*ok411*)] created in our lab.

All nematodes (hereafter referred to as worms) used in experiments were synchronized at the L1 stage using a bleach/sodium hydroxide mixture optimized following standard techniques (Porta-de-la-Riva et al. 2012). Worms reared in liquid culture were maintained on a shaker set to 120 rpm, at 20° C. Liquid cultures were maintained in S-complete media and fed using the UV sensitive strain NEC937, following UV exposure per N. Stroustrup (CRG, Barcelona, personal communication).

### Genotyping

Strains sourced from the CGC and those resulting from an in-house cross were genotyped using standard polymerase chain reaction (PCR) techniques. Primers used to check the *cat-1* (*ok411*) mutant allele are found in the description listing for strain RB681 on the website for the CGC (Caenorhabditis_Genetics_Center 2020).

### Western Blot

To visualize the *cat-1*/VMAT protein, *C. elegans* strains N2, *cat-1* (*ok411*), *cat-1* (*e1111*), and *dat-1* (*ok157*) were cultured on NGM plates. 50 worms of each strain were transferred to a 1.5ml Eppendorf tube containing 150 μl of M9. After washing three times with M9, the worm pellet was suspended in 25 μl of M9, snap frozen in liquid nitrogen, and stored at −80°C until use. Prior to analysis, samples were thawed on ice, after which 25μl of Laemmli 2× sample buffer and proteinase inhibitor cocktail were added and the sample mixtures were placed in a sonicating water bath at room temperature for 2 minutes and immediately placed in heat block at 95°C for 5 minutes. Samples were run on a NuPage 10% bis tris gel (Thermo Fisher, (Waltham, MA)) and transferred to a PVDF membrane. Nonspecific antibody binding was blocked with a 7.5% milk in tris buffered saline plus tween (TBST) solution for 1.5 hours at room temperature. The primary antibody used was a polyclonal rabbit anti *cat-1*/VMAT serum generated by Dr. Richard Nass (IUSM, Indiana) at a dilution of 1:1,000. The Secondary antibody used was a goat anti-rabbit HRP-conjugated at a dilution of 1:5,000) (Jackson ImmunoResearch (West Grove, PA)).

### Immunohistochemistry

The method was based on a previously published Freeze-Cracking protocol unless otherwise specified (Duerr 2013). Poly-l-lysine coated slides were prepared to bind worms to slides. Approximately 5,000 worms per strain were used to prepare multiple slides. A Fisherbrand™ Cooling Cartridge (Thermo Fisher; Waltham, MA) pre-chilled overnight in −80°C was used to freeze the slides. The slides were lightly fixed using the methanol-acetone method. Nonspecific antibody binding was blocked with a 10% Goat Serum (Thermo Fisher (Waltham, MA)) overnight at 4°C. The primary antibody used was a polyclonal rabbit anti *cat-1*/VMAT serum generated by Dr. Richard Nass (IUSM, Indiana) at a dilution of 1:100. The secondary antibody used was Goat anti-Rabbit IgG (H+L) Cross-Adsorbed Secondary Antibody, Alexa Fluor 488 (Thermo Fisher; Waltham, MA). VECTASHIELD Antifade mounting medium with DAPI (Vector Laboratories; Burlingame, CA) was added prior to sealing the coverslips with nail polish. Images were taken at 40x magnification on a Leica DMi8 (Leica; Wetzlar, Germany) inverted epifluorescent microscope.

### Analysis of monoamine derived behaviors

To create reserpine assay plates, reserpine was dissolved in glacial acetic acid (50mM) and diluted 1:80 in M9 for a final concentration of 625μM. 400μl was added to the treatment plate and allowed to dry before transferring worms (Duerr et al. 1999). Worms were seeded onto growth plates (6 cm NGM with live OP50) as synchronized L1s and allowed to grow for 48 hours. Worms were transferred to reserpine plates and left for 12 hours prior to behavioral assays except for egg laying where worms were kept on reserpine plates until the second day of adulthood.

#### Grazing

Plates for assessing grazing behaviors were created per Duerr *et. al*. (Duerr et al. 1999). For our assays, worms were rinsed off treatment plates and spotted in 20μl of M9 with approximately 100 worms per assay. A lint free tissue was used to wick away the M9 and the worms were recorded using a FLIR chameleon 3 camera from Edmund Optics (Barrington, NJ) until either the last worm entered the lawn or a maximum of 30 minutes. Videos were scored by a researcher blind to treatment conditions, measuring the amount of time it took for each worm to enter the lawn of OP50 (tip of nose to end of tail).

#### Pharyngeal Pumping

Pharyngeal contractions were scored by observation of worms through a standard stereomicroscope, using finger taps on an electronic counting application while a timer was running, a method similar to that previously described (Miller et al. 1996).

#### Wave Initiation Rate

The celeST software package was used as described to determine aspects of swim behavior for the N2, *cat-1* (*ok411*) and N2 worms treated with reserpine groups (Restif et al. 2014). Briefly, 4 worms were placed in 60 ul of M9 on a glass slide. Recordings of swim behavior were made as a series of jpeg images using a chameleon 3 camera ((FLIR) Wilsonville, OR)) for 30s at a frame rate of 18 f/s. All subsequent analysis was completely automated using the celeST software package.

#### Egg laving

Retention of eggs inside worms was determined by dissolving 2-day-old adult worms grown on agar in 20% hypochlorite in a 96 well plate and counting the number of eggs released.

### MPP^+^ Treatment

Synchronized worms were grown using standard liquid culture (see above) using 125 ml erlenmeyer flasks until they reached the YA stage of development. At this point, worms were sorted using confirmed gating parameters into 96 well plates using the COPAS FP-250 large particle flow cytometer (Union Biometrica, MA) to select for fully developed YA worms. Various concentrations of MPP^+^ dissolved in water were added to the liquid culture (s-complete, 100uM Floxuridine), and the worms were left at 20°C on a shaker for an additional 48 hours. Worms were then anesthetized in 10mM sodium azide and the area surrounding the 6 dopamine neurons in the head were photographed at 20x magnification using a Fisher Scientific EVOS epifluorescence microscope (Waltham, MA) for later analysis. All photographs were taken using identical objective, light intensity, and exposure settings. We used the Image J software to determine neurodegeneration based on the total area of fluorescence post 48h of treatment, with the observation that worms treated with MPP^+^ exhibit overall reductions of dopamine neuron size and branching (Schindelin et al. 2012). First, photographs were converted into 8-bit grayscale tiff files. Following conversion, the photographs were batch processed using the Yen automated thresholding algorithm (Yen et al. 1995). Following thresholding, total area of GFP fluorescence was calculated using the particle analysis plugin. Results were combined over 4 experiments as percent area of untreated worms. Statistical testing using one-way ANOVA followed by Tukey HSD multiple comparisons were designed to compare the extent of degeneration between *pDAT*::GFP worms and *pDAT*::GFP worms lacking the *cat-1*/VMAT protein (*pDAT*::GFP; *cat-1* (*ok411*)) when treated with equivalent doses of MPP^+^. A small series of treatments were run through the COPAS biosorter to get representative fluorescent profiles of worms treated with MPP^+^.

### High resolution mass spectrometry-based metabolomics

We applied untargeted high resolution mass spectrometry-based metabolomics (HRM) to characterize metabolic effects of the *cat-1* mutation and MPP^+^ exposure in N2 worms. Worms with the *cat-1* (*ok411*) mutation and wildtype N2 worms were grown in liquid culture as described above and 6 replicates per strain were collected in M9 at the L4 stage and snap frozen in liquid nitrogen. Prior to MPP^+^ exposure, N2 worms were grown in liquid culture and at the L4 stage, 500 worms were sorted into wells of a 24-well plate using the COPAS FP-250. Worms were exposed to 1mM MPP^+^ or the control for 4 hours, washed and snap frozen. All samples were stored at −80°C until processed. Metabolites were extracted using acetonitrile (in a 2:1 ratio) which was added to samples along with an internal standard (Soltow et al. 2013). Each sample was placed in the bead beater to disrupt the cuticle, and included shaking at 6.5 m/s for 30 seconds, cooling on ice for one minute, and placed in the beater for another 30 seconds, at the same speed (Mor 2020). All processing was performed on ice or in a cold room when necessary. Untargeted high-resolution mass spectrometry analysis was performed using a dual-chromatography and acetonitrile gradient that included HILIC chromatography with positive electrospray ionization (HIILIC-pos) and C18 column with negative electrospray ionization (C18-neg) (Walker et al. 2019). Mass spectral data was generated on a ThermoFischer Q-Exactive HF Orbitrap mass spectrometer operated at 120,000 resolution over mass-to-charge (*m/z*) scan range 85 to 1250. Data were extracted using the R packages apLCMS (Yu et al. 2009) and xMSanalyzer (Uppal et al. 2013).

The HILIC positive column measured 21,479 uniquely detected mass spectral features, identified by accurate *m/z*, retention time and intensity in each sample. To focus on detection of endogenous metabolites and pathways related to neurotransmitter metabolism, data analysis was restricted to the HILIC-pos condition only.

### *cat-1*/VMAT and MPP^+^ metabolome wide association study

We analyzed HRM results using three separate approaches to evaluate metabolic alterations due to the *cat-1* mutation and MPP^+^ exposure: 1) The metabolic effect of the *cat-1* mutation was characterized by comparison to wildtype N2 worms grown under identical conditions. 2) Disruption to worm metabolic function following MPP^+^ exposure for exposed N2 worms and untreated N2 controls and 3) An overlap analysis was completed by identifying features associated with both *cat-1* and MPP^+^ for similarities in disruption of systemic metabolic response due to altered dopaminergic neuronal health. A feature was retained in each of the first two analyses if its abundance in at least one worm sample was 1.5 times its abundance in the M9 buffer used to wash and collect the worms. Prior to data analysis, missing values for each feature were imputed with half the value of the minimum abundance, quantile normalized, log10 transformed, and auto scaled. Features were analyzed using t-tests, partial least squares discriminant analysis (PLS-DA), hierarchical clustering, and pathway analysis using mummichog (Li et al. 2013). All data processing, analysis and visualization was done in R version 3.6.0, using functions: preprocessCore::normalize.quantiles() (Bolstad et al. 2003), (RFmarkerDetector::autoscale() (Palla 2015), gplots::heatmap2() (Gregory R. Warnes 2019), ggplot2::ggplot() (Wickham 2016), and mixOmics::plsda() (Kim-Anh Le Cao 2016). Pathway analysis was completed using the Mummichog algorithm (Li et al. 2013) hosted on the MetaboAnalayst (www.metaboanalyst.ca) module “MS peaks to Pathway” (Chong et al. 2019), using the *Caenorhabditis elegans* metabolic reference map available through KEGG. Using output from the t-tests, a nominal p-value cut-off of 0.1 was used to select features for pathway analysis, using a mass tolerance of 5ppm for *cat-1* (*ok411*) mutant and for wild-type worms treated with MPP^+^. Features that were different between wildtype and *cat-1*(*ok411*) worms at p < 0.1 (356 features) were compared to features that were different between MPP^+^ treated and control worms at p < 0.1 (801 features). To discover overlapping features, we used the *getVenn()* function in the xMSanalyzer R package with a mass-to-charge tolerance of 5 ppm and retention time tolerance of 5 seconds. We used the xMSanalyzer::feat.batch.annotation.KEGG() function to determine level 4 annotations, (Schymanski rating ((Schymanski et al. 2014)) for the overlapping features (Uppal et al. 2013).

### Statistics

All data excluding metabolomics were analyzed using the Graphpad statistical software package (San Diego, CA). All metabolomic analyses were conducted in R (version 4.0.2).

## Results

### Phylogenetic relationships of vesicular amine transporters

Cladistic relationships amongst vertebrate and invertebrate species between the vesicular monoamine transporter(s) and the vesicular acetylcholine transporter are shown in Figure 1.

**Figure 1.**
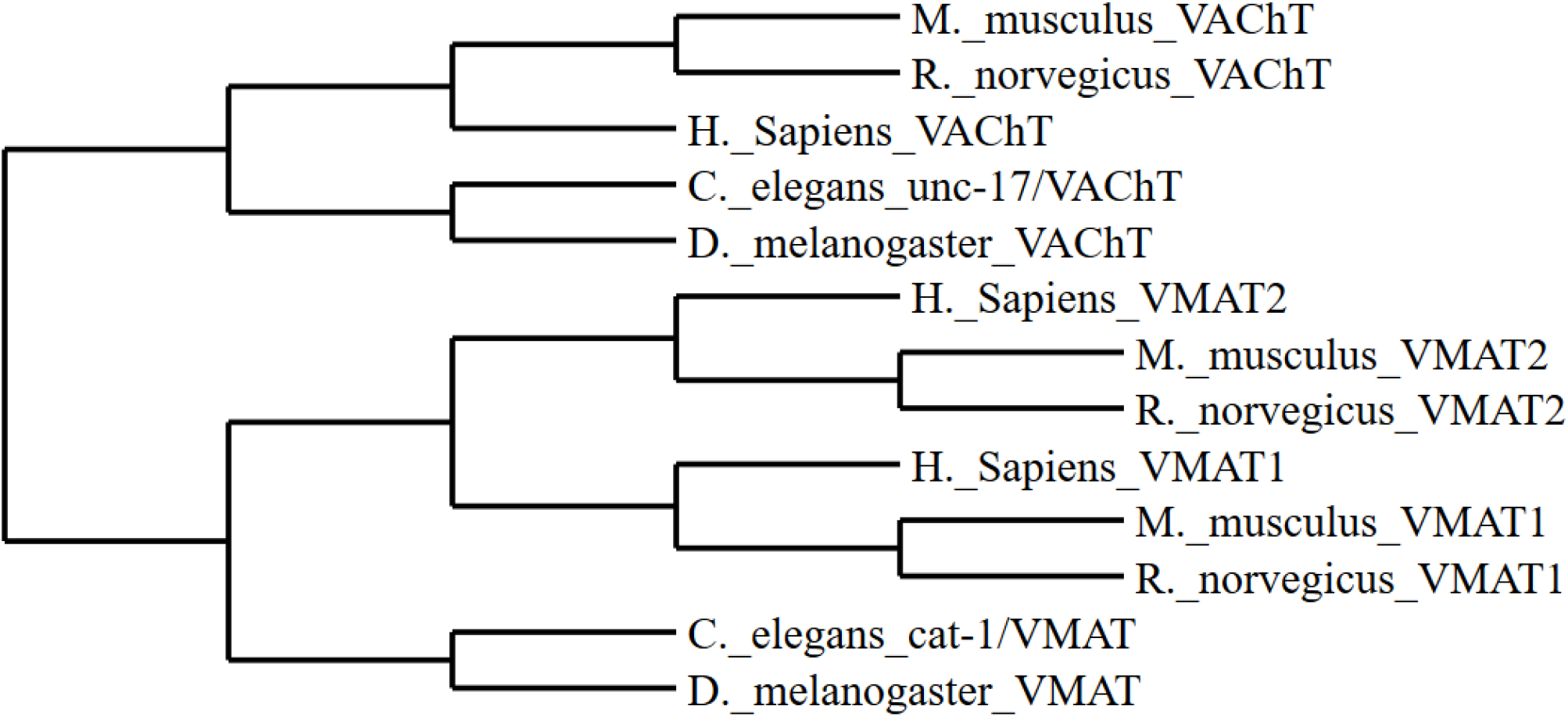
Cladogram representing species specific relationships amongst vesicular amine transporters

### Genotyping and protein analysis of *cat-1* (*ok411*)

The *ok411* mutant allele was produced by the high throughput *C. elegans* gene knockout consortium (Consortium 2012). The mutation is a 429 basepair deletion resulting in complete loss of the first coding exon in the *cat-1* gene and a loss of the *cat-1*/VMAT protein by western blot and immunofluoresence (Figure 2, A, B, C). A breeding cross of the BZ555 *pDAT::GFP* with the *cat-1*(*ok411*) mutant was successful in producing a transgenic with the mutant *cat-1*(*ok411*) allele (Figure 2, A). Our efforts at immunostaining support the work of previous labs indicating the expression pattern of *cat-1*/VMAT (Duerr et al. 1999; Serrano-Saiz et al. 2017).

**Figure 2.**
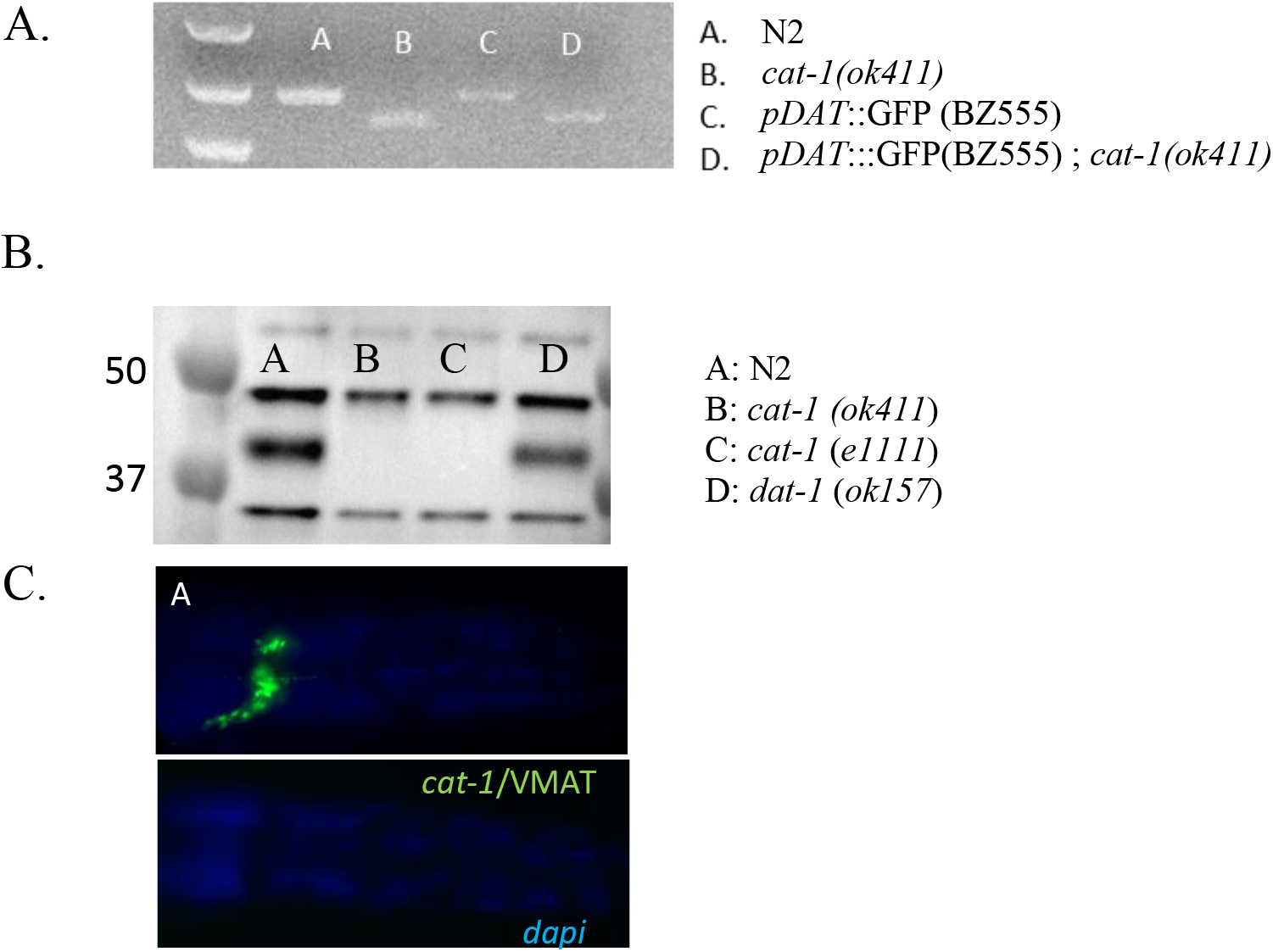
The *cat-1* (*ok411*) strain is deficient in the production of cat-1/VMAT protein. **A)** Primers specific for *cat-1* reveal a 400 bp deletion in the *cat-1* (*ok411*) mutant strain **B)** The strain *cat-1* (*ok411*) does not produce *cat-1*/VMAT protein. **C)** The *cat-1*(*ok411*) strain shows no evidence of *cat-1*/VMAT immunoreactivity (DAPI staining in blue showing outline of worm).

### Monoamine derived behaviors

The *cat-1*(*ok411*) mutant strain lacking *cat-1*/VMAT protein and pharmacological inhibition of *cat-1*/VMAT in N2 worms treated with reserpine demonstrated defective grazing behavior (Figure 3, A) indicative of disruption of dopamine and serotonin signaling. In order to determine if this behavior is due to a general difference in motility, we subjected the *cat-1*(*ok411*) strain and the reserpine treated N2 worms to automated swim behavior analysis using the celeST software and found no significant difference between any of the groups using 10 different measures of stroke analysis including a measure (wave initiation rate measured as: waves / minute) analogous to the thrashing assay (Figure 3, D). We employed two measures of serotonergic behaviors; egg laying and pharyngeal pumping. Worms with disrupted serotonergic signaling tend to hold their eggs *in-utero* and have a reduced rate of pharyngeal pumping. We found the *cat-1*(*ok411*) strain and N2 wild type worms treated with reserpine to be both deficient in egg laying behavior and pharyngeal pumping (Figure 3, B & C).

**Figure 3.**
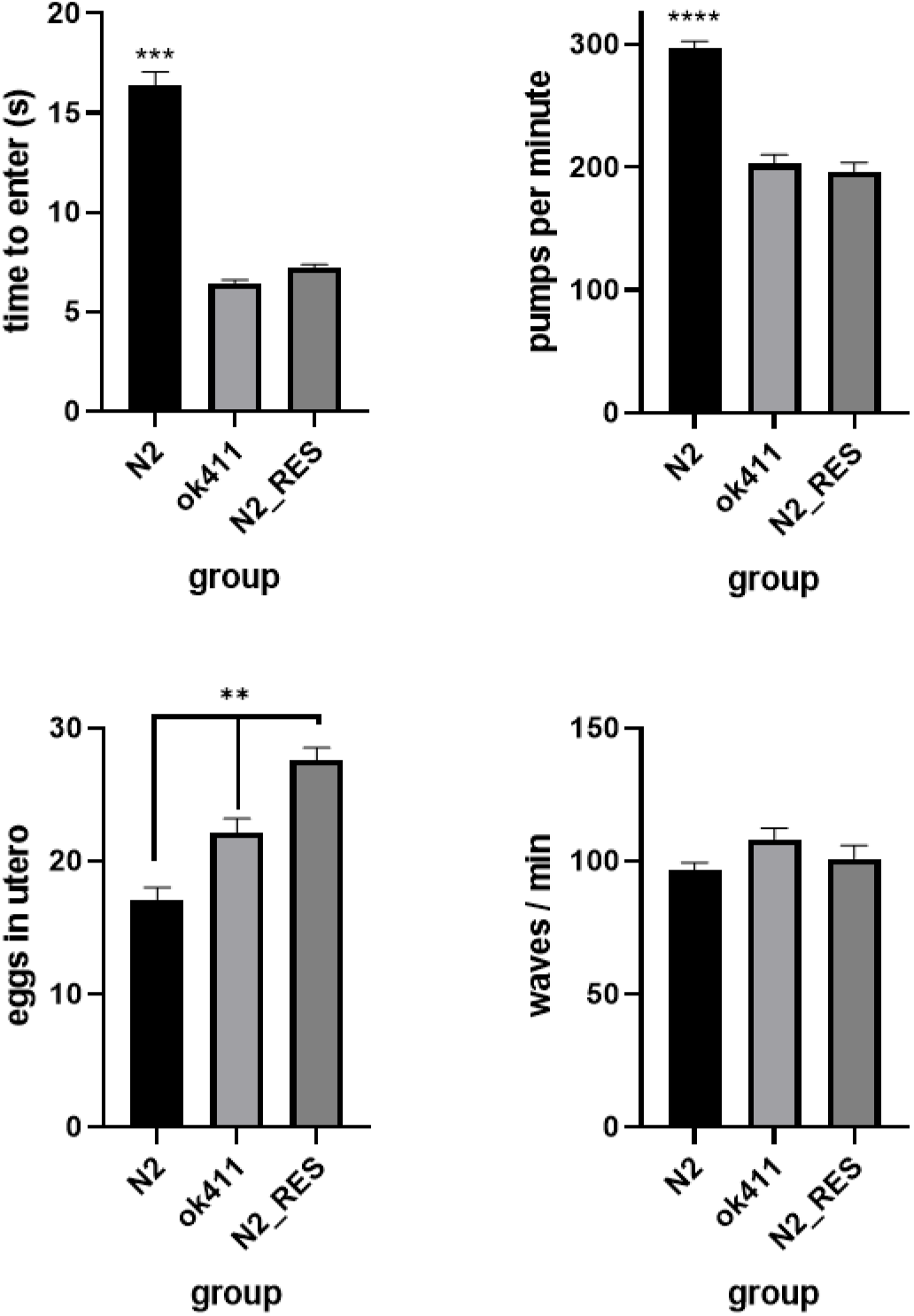
The loss of *cat-1*/VMAT protein results in altered monoaminergic behaviors. **A)** The grazing response is deficient in the *cat-1* (*ok411*) mutant strain and N2 worms treated with reserpine. **B)** *cat-1* (*ok411*) worms and N2 worms treated with reserpine have impaired pharyngeal pumping compared to N2. **C)** *cat-1* (*ok411*) and N2 worms treated with reserpine are deficient in egg laying (more eggs in utero). **D)** There is no difference in wave initiation between N2 and the *cat-1* (*ok411*) mutant. ** *p* < .01, *** *p* < .001 vs. all other groups, **** *p* < .0001 vs. all other groups

### MPP^+^ exposure and neurodegeneration

Worms treated with MPP^+^ exhibit reductions in total area consistent with the degeneration of dopamine neurons (Figure 4, B). Representative fluorescent profiles using the COPAS biosorter support this assertion (Figure 4, C). We find that BZ555 transgenic worms lacking *cat-1*/VMAT protein are more susceptible to the toxic effects of MPP^+^ than the BZ555 transgenic worms with *cat-1*/VMAT protein (Figure 4, A). Reductions in total GFP area occur at lower concentrations of MPP^+^ in worms with the *cat-1*(*ok411*) mutant allele.

**Figure 4.**
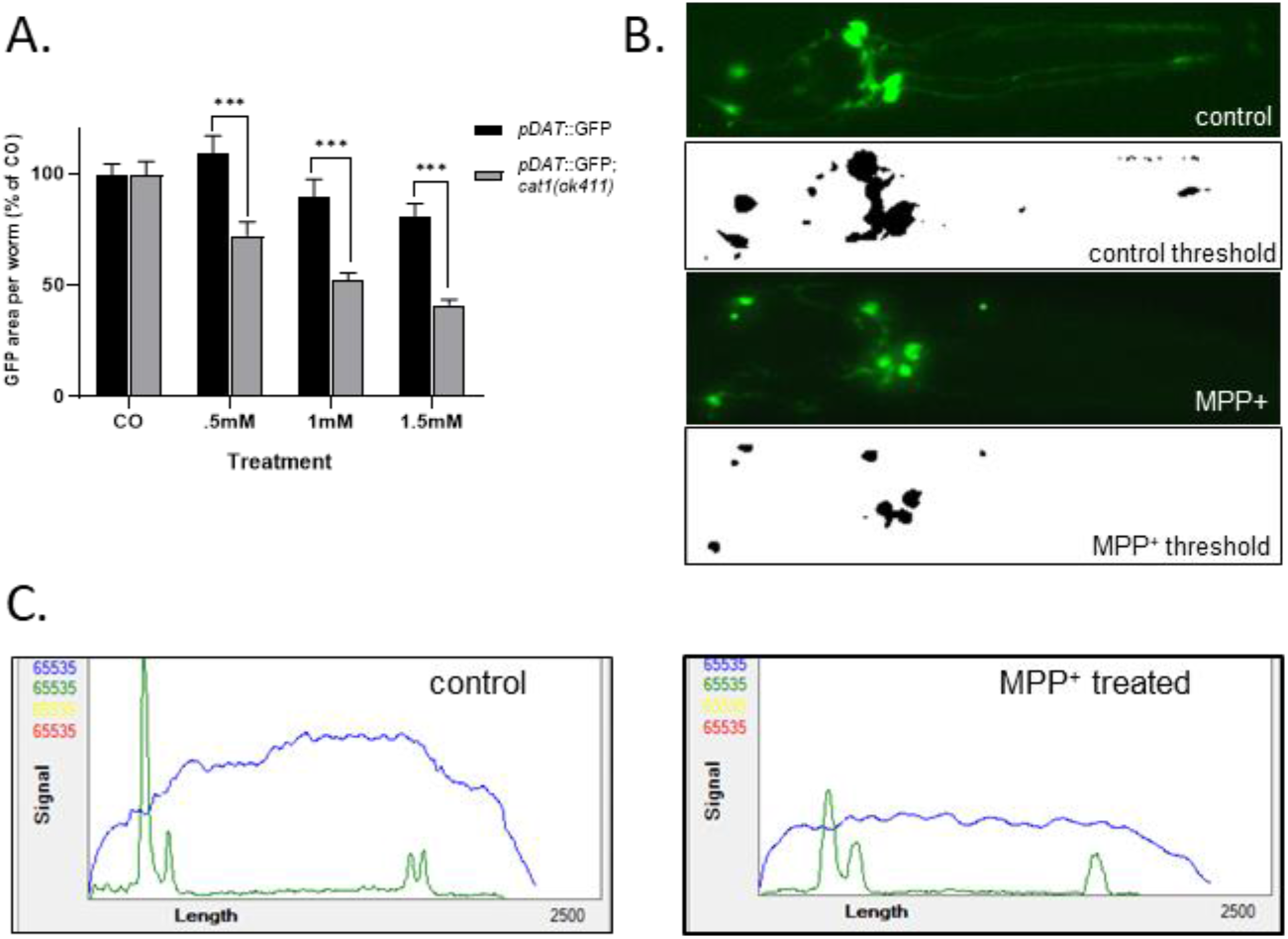
Dopamine neurons lacking the *cat-1*/VMAT transporter are more susceptible to MPP^+^ induced neurodegeneration. Image analysis using automated thresholding shows increased sensitivity of the *cat-1*/VMAT mutant *ok411* to MPP^+^ via reduced fluorescent area of dopamine neurons (A & B). C. Representative profile graph using the COPAS large particle flow cytometer. GFP fluorescence driven by the *dat-1* promoter. \*\*\**p* < .0001

### Metabolomics

There were 114 features (in red) different between the two groups with p< 0.05 (Figure 5, A). There was also clear clustering of the features with p < 0.05 between the two groups (Figure 5, B). A metabolome wide association study of phenylalanine metabolism has shown that for initial discovery purposes, raw p combined with metabolic pathway enrichment improves detection of biological effect while reducing identification of false positive biological effects (Go et al. 2015). Thus, for metabolic pathway enrichment, we considered all 356 features with p < 0.1. It revealed several metabolic pathways altered with a fisher exact test p-value < 0.1 in the *cat-1*(*ok411*) mutant (Figure 5, C), including tyrosine metabolism (Figure 5, C). A partial least squares differential analysis showed separation of the wild type N2 and *cat-1*(*ok411*) groups (Figure 5, D).

**Figure 5.**
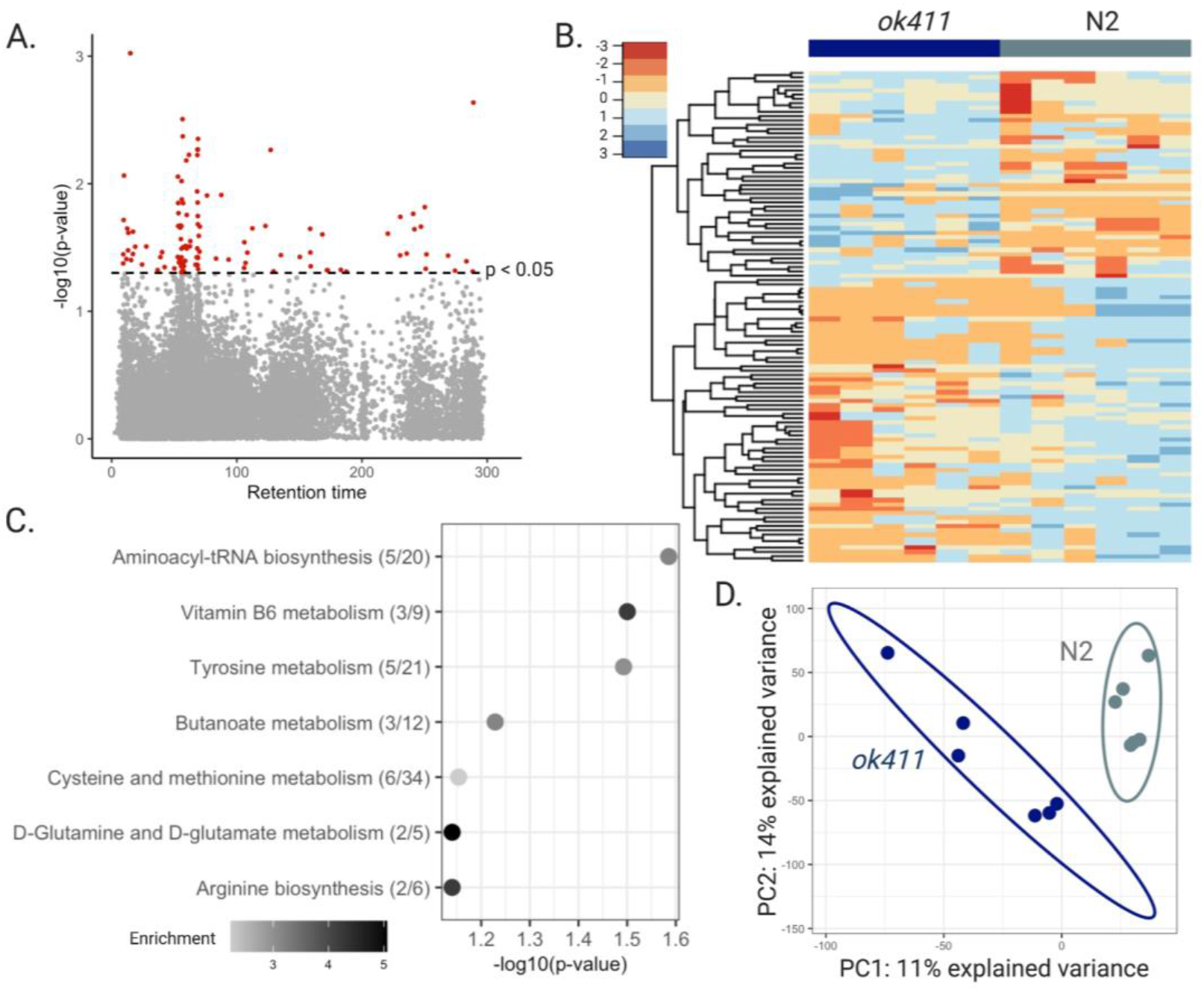
The *cat-1* (*ok411*) strain shows patterns of altered metabolism. **A)** Manhattan plot shows features that were different with *p* < 0.05 (red) in *cat-1* (*ok411*) mutants compared to wild type N2 worms, **B)** Hierarchical clustering of features associated with the *cat-1* (*ok411*) mutant with p < 0.05, **C)** Top pathways altered (fisher exact test p-value < 0.1) in *cat-1* (*ok411*) worms, analyzed using the Mummichog software. The overlap size of the pathway is indicated in parentheses (number of significant hits/pathway size). The color of the bubbles represent enrichment, calculated as the quotient of total number of hits in the pathway divided by expected number of hits. **D)** Partial least squares discriminant analysis (PLS-DA) comparing *cat-1* (*ok411*) mutants to wild type N2 worms.

Untargeted metabolomic analysis of worms exposed to MPP^+^ for 4 hours was performed similarly to the *cat-1*(*ok411*) mutant. There were 199 features (in red) significantly different between the treated and untreated groups with *p*< 0.05 (Figure 6, A). These significant features clustered differently in the two groups as seen in the heatmap generated using hierarchical clustering analysis (Figure 6, B). Pathway analysis using features with *p* < 0.1 (801 features) was done using mummichog hosted on MetaboAnalyst. It revealed several pathways of interest (Figure 6, C), including the tyrosine and tryptophan metabolism pathway, glycerophospholipid metabolism and the pentose glucuronate interconversions pathways. (Figure 6, C). A partial least squares discriminant analysis was able to differentiate between the treatment groups (Figure 6, D). Comparing features with *p* < 0.1 in *cat-1*(*ok411*) worms and worms exposed to MPP^+^ revealed 14 features altered in both conditions (Supplemental figure 1. A). KEGG annotations were made for 5 of the 14 features (Supplemental table 1) with level 4 confidence according to Schymanski criteria (Schymanski et al. 2014). See supplemental excel workbook for raw data, list of all pathways enriched, and the variable importance score of features analyzed using PLS-DA.

**Figure 6.**
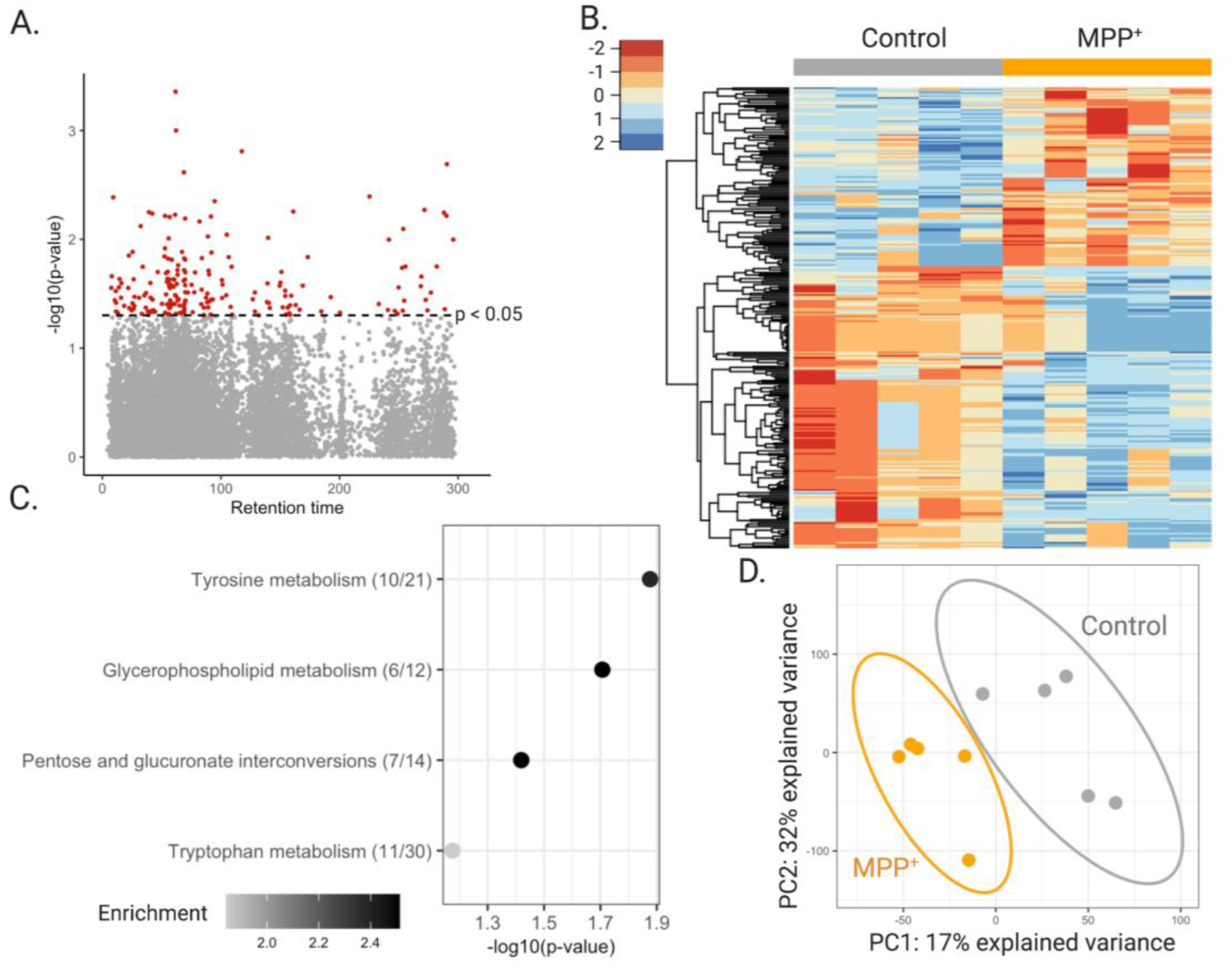
Worms treated with MPP^+^ show patterns of altered metabolism. **A)** Manhattan plot shows features that were different with p < 0.05 (red) N2 worms treated with MPP^+^ compared to untreated N2 worms, **B)** Hierarchical clustering of features associated with treatment with *p* < 0.05, **C)** Top pathways altered (fisher’s exact test p-value < 0.1) in N2 worms treated with 1mM MPP^+^, analyzed using the Mummichog software. The overlap size of the pathway is indicated in parentheses (number of significant hits/pathway size). The color of the bubbles represent enrichment, calculated as the quotient of total number of hits in the pathway divided by expected number of hits, **D)** Partial least squares discriminant analysis (PLS-DA) comparing N2 worms with N2 worms treated with 1mM MPP^+^.

## Discussion

The importance of vesicular amine transporters in mitigating catecholaminergic toxicity is well established (Guillot and Miller 2009; Lohr and Miller 2014; Spina and Cohen 1989). Multiple *in vivo* studies have demonstrated the toxic consequences of dopamine including early work that demonstrated toxicity resulting from striatal injections of exogenous dopamine (Hastings et al. 1996). Later studies investigated the consequences of dysregulating endogenous dopamine handling. Overexpression of the plasmalemmal dopamine transporter (DAT) resulted in increased dopamine metabolism and nigrostriatal degeneration thought to result from an accumulation in cytosolic dopamine with insufficient vesicular sequestration capabilities (Masoud et al. 2015). Furthermore, introducing DAT into striatal interneurons that lack the proper machinery to sequester dopamine within vesicles resulted in neurodegeneration, demonstrating the necessity of proper dopaminergic handling for neuronal health (Chen et al. 2008). Mice deficient in VMAT2 display indices of cytosolic dopamine metabolism as well as age-dependent nigrostriatal dopaminergic degeneration (Caudle et al. 2007; Taylor et al. 2014; Taylor et al. 2009). This work has been replicated in *Drosophila melanogaster* deficient in VMAT expression that display fewer dopaminergic neurons (Lawal et al. 2010). The purpose of this paper is to establish the *cat-1* knockout specifically and the *C. elegans* model in general as a viable model for the role of vesicular monoamine transporter as a mediator of synaptic function and neuronal vulnerability.

Single worm PCR and the utility of the hermaphroditic reproduction strategy of these worms allows us to readily identify mutants and perform relevant crosses. Simple breeding schemes allow us to create enough reporter strains to examine the effects of vesicular transporter disruption for the entire *C. elegans* neural network. Western blot and immunofluorescent techniques, though rarely used in *C. elegans* research, provide a valuable way of confirming the presence or absence of a protein and subsequent distribution in relevant tissues that are consistent with methods used in mammalian models.

Despite their many advantages, studies of *C. elegans* focused on the *cat-1*/VMAT protein are limited (Duerr et al. 1999; Young et al. 2018). The importance of *cat-1*/VMAT in the establishment of food detection, feeding rate and reproduction have been demonstrated in the past and replicated by the genetic and pharmacological interventions used in this study (Duerr et al. 1999; Young et al. 2018). Previous cell ablation studies demonstrate that the slowing down of a worm when on a food source is dependent on the presence of intact dopamine neuronal architecture (Sawin et al. 2000). It is interesting that the loss of functional *cat-1*/VMAT replicates this phenotype with the probable effect of system wide catecholaminergic dysfunction (Figure 3, A and (Duerr et al. 1999; Young et al. 2018)). Indeed, one other group has found the addition of either exogenous serotonin or pramipexole can rescue this behavior in *cat-1*/VMAT mutant strains (Young et al. 2018). Although beyond the scope of this paper, further exploration of this behavior may reveal synergistic or antagonistic patterns of catecholamine function. Optimal rates of pharyngeal pumping and egg laying require the input of serotonin (Avery and Horvitz 1990; Trent et al. 1983). The deficits observed in pharyngeal pumping and egg laying rate seen in the *cat-1(ok411)* mutant and associated pharmacological treatment with the inhibitor reserpine suggest that proper storage and trafficking of endogenous stores of this catecholamine is important for its function.

The synaptic vesicle containing functional VMAT2 provides protection to the cell from both endogenous (oxidized dopamine) and certain exogenous (MPTP) insults (Guillot and Miller 2009; Spina and Cohen 1989). The MPTP model of selective toxicity to dopamine neurons is well established and a hallmark of mouse work in the PD field (Meredith and Rademacher 2011). The *C. elegans* model has been used many times to demonstrate the selective toxicity of MPP^+^ (the active metabolite of MPTP) on dopaminergic architecture, however, to our knowledge no laboratory has tested the effects of MPP^+^ in worms lacking the *cat-1*/VMAT protein (Braungart et al. 2004; Lu et al. 2010; Pu and Le 2008; Wang et al. 2007; Yao et al. 2011). These findings suggest what we have previously found in mice, namely that levels of VMAT regulate MPTP vulnerability (Lohr et al. 2016). Given the genetic tractability of *C. elegans*, the creation of *cat-1*/VMAT overexpressing worms in future studies is certainly feasible and may facilitate future studies aimed at using enhanced monoamine storage as a therapeutic intervention. It is notable, but not surprising given the evolutionary development of the TEXAN antiporters that the protective effects of monoamine transporters extends across both vertebrate and invertebrate animal models (Figure 1; (Schuldiner et al. 1995)). This provides further confidence in the utilization of *C. elegans* as a model for vesicular dysfunction related to monoamine transporter efficacy.

*C. elegans* is gaining support as a model for toxicity testing, demonstrating conserved LD50 values and similarities in toxic mechanisms with rodent models (Hunt 2017). Previous research demonstrates a clear connection between a number of environmental pollutants and the expression of VMAT2 in the nigrostriatal system of mouse models. However, mechanistic studies and full-scale screens of Toxcast libraries are lacking due to the absence of an inexpensive, viable model of monoaminergic vesicular dysfunction. The introduction of the COPAS large particle cytometer may help provide the resolution needed to bring high throughput capability to the screening of potential neurotoxic compounds. Indeed, we are currently refining assays in our lab (Figure 4, C). The characterization of the *C. elegans* model of impaired vesicular storage seen in this paper is a promising development.

Metabolomics analysis of *cat-1*(*ok411*) worms showed altered pathways in tyrosine metabolism, possibly resulting from the vesicular mishandling of dopamine in vesicles lacking *cat-1*/VMAT. Members of the pathway suggested lower dopamine levels and higher levels of dopamine metabolites (supplemental Figure 3, A). Pathway analysis also suggested changes in glutamate metabolism in worms lacking *cat-1*/VMAT, similar to a previous study that showed altered levels of glutamate detected through NMR metabolomics in the substantia nigra of aged mice expressing low levels of VMAT2 (Salek et al. 2008). MPP^+^ enters the neuron through the cell membrane via dopamine transporter 1 (DAT1), thus selectively affecting dopaminergic neurons (Gainetdinov et al. 1997). As such, it was telling to see differences in the metabolism of tyrosine − the precursor of dopamine – in wildtype worms treated with MPP^+^. The finding of differences in tryptophan metabolism is more curious considering that MPP^+^ has low affinity for the plasma membrane transporter in serotonin neurons. However, cellular crisis in dopaminergic cells could lead to alterations in tryptophan metabolism reflecting the interdependence of both amines. In fact, these changes in tryptophan metabolism might represent a more generalized response to the increased proteotoxicity involved in cellular degeneration (van der Goot et al. 2012). Likewise, changes in the pentose glucuronate interconversion pathway may also be indicative of a mitochondrial crisis (Mullarky and Cantley 2015). Features that are altered in both types of dopaminergic neuronal perturbations could provide information on metabolic pathways that are of interest to neuronal damage and degeneration. While we were able to discover 14 features overlapping between the two conditions, we did not have sufficient power to determine pathways of interest that were altered between the two conditions. Future studies could use spectral fragmentation to identify the chemical structure of these 14 features. Further, chemical entities of metabolic relevance can be studied by using an appropriate mutant worm strain or by employing biochemical assays.

High resolution mass spectrometry-based metabolomics has been used in several *C. elegans* studies (Edison et al. 2015; Hastings et al. 2017; Mor 2020; Witting et al. 2018). The comprehensive molecular understanding of the nematode model makes it amenable to characterizing systemic biochemistry and perturbations to global metabolism as a result of genetic, environmental and toxicological perturbations. Our initial findings suggest that the use of this technique could add to the systems-level evaluation of toxicity in this model. This information may provide further avenues of exploration when looking at metabolic effects of neurodegeneration. In fact, systems-wide approaches to studying common neurodegenerative diseases like PD are recommended by the variety of circuitry affected (Taylor et al. 2014; Taylor et al. 2011; Taylor et al. 2009). For example, in the few cases of dopamine-serotonin transport disease, a disease related to a mutant allele in VMAT2 known as P387L, patients exhibit developmental delay, parkinsonism, sleep disturbances, mood disorders and a host of issues related to autonomic dysfunction (Rath et al. 2017). The application of high resolution mass spectrometry allows us to identify metabolic dysfunction in an untargeted manner. The unbiased nature of this approach has the capability of providing links to associated human diseases previously unrecognized. By replicating the dysfunction of monoamine transport and storage in *C. elegans* we may yet uncover similar metabolic signatures of relevant disease in humans. Our group has recently reported that the same high resolution mass spectrometry-based metabolomic platform can identify disease-specific alterations in plasma from patients with and without Alzheimer’s disease (Niedzwiecki et al. 2019; Vardarajan et al. 2020). demonstrating the utility of the approach from worms to humans. We expect *C. elegans* will play an important role in future exposome-level studies by providing a laboratory-based validation of observed metabolomic alterations in human disease (Vermeulen et al. 2020). Overlap in common metabolic signatures could potentially be useful for treatment and early diagnosis in neurodegenerative diseases and a number of addiction and mood disorders.

In summary, our results demonstrate the evolutionary-conserved nature of monoamine function in *C. elegans* and further suggest that high-resolution mass spectrometry-based metabolomics can be used in this model to study environmental and genetic contributors to complex human disease.

## Funding

This work was supported by the National Institute of Environmental Health Sciences (R01ES023839, P30ES019776, P30ES009089, U2CES02656, T32 ES012870).

## Acknowledgements

We would like to acknowledge several members of the *C. elegans* community for their kind assistance. The Katz lab at Emory University, especially Teresa Lee who got us started in the basics of the *C. elegans* model system. Nicholas Stroustrup and Megan Niedzwiecki for assistance with establishing the Lifespan machine in our lab. Richard Nass for providing several antibodies, including the *cat-1*/VMAT antibody used in this paper. The Edison lab for their hospitality and discussions of worm metabolomics. The Blakely lab for helpful discussions on metabolism in *C. elegans*. Finally, we would like to thank Shuzhao Li for help with Mummichog pathway analysis, and ViLinh Tran and Michael Orr from the Clinical Biomarkers lab at Emory University for running our metabolomics samples and creating biochemical assays for redox analysis in worms.

## Notes

### Competing Interest Statement

The authors have declared no competing interest.

